# Comparison of different computational frameworks for metabolic modeling from single-cell transcriptomics data in glioblastoma

**DOI:** 10.64898/2026.02.28.706749

**Authors:** Mirte De Temmerman, Boris Vandemoortele, Vermeirssen Vanessa

## Abstract

Metabolic reprogramming is a hallmark of glioblastoma, yet how distinct malignant and tumor microenvironment cell populations contribute to this metabolic heterogeneity remains poorly defined. Since direct single-cell metabolomics remains technically limited, transcriptomics-based computational inference offers a powerful alternative. Here we apply and systematically compare three complementary computational methods: (1) metabolic pathway activity scoring, (2) gene regulatory network inference focused on metabolic enzyme gene regulation, and (3) single-cell metabolic flux prediction. These methods were applied to snRNA-seq data from a set of GBM patient samples using the Human1 genome-scale metabolic model as a unified reaction and pathway annotation prior knowledge reference. Across all three methods, tumor-associated macrophages emerge as the metabolically dominant tumor microenvironment population. Tumor-associated macrophages in mesenchymal-like tumors show coordinated transcriptional control of lipid metabolism by five recurrently active transcription factors. They also exhibit consistent nucleotide biosynthesis flux and glutamate-to-glutamine conversion potentially supporting malignant cells. These findings demonstrate that multi-layered metabolic inference can resolve cell-type/state-specific dependencies in glioblastoma and highlight tumor-associated macrophage metabolism as a promising therapeutic target

## Introduction

Glioblastoma (GBM) remains the most aggressive adult primary brain malignancy and continues to pose major therapeutic challenges despite advances in molecular profiling and multimodal treatment strategies. Standard-of-care therapy includes maximal safe resection followed by radiotherapy and chemotherapy with temozolomide, but this only yields a median survival of 14 to 18 months and essentially all tumors recur^1,2^. Over the past decade, large-scale genomic and epigenomic studies have provided detailed maps of inter-patient heterogeneity, leading to the classification of IDH-wild-type GBM into proneural (PN-like), classical (CL-like), and mesenchymal (MES-like) molecular subtypes^3,4^. However, targeted therapies informed by bulk tumor subtypes have largely failed in clinical trials, highlighting the importance of intratumoral heterogeneity, which cannot be resolved at bulk resolution^5–7^.

Single-cell RNA sequencing (scRNA-seq) has transformed our understanding of GBM biology by revealing that each tumor comprises a dynamic mixture of malignant cell states that map onto neurodevelopmental lineages, including neural progenitor-like (NPC-like), oligodendrocyte progenitor-like (OPC-like), astrocyte-like (AC-like), and mesenchymal-like (MES-like) states^8–11^. Cells continuously transition between states in response to microenvironmental cues and therapy, creating heterogeneous and plastic populations with distinct vulnerabilities. Recent studies integrating copy-number profiles, lineage trajectories, and regulon activity maps have demonstrated that these cell states are shaped by oncogenic programs and tumor microenvironment (TME) interactions^12,11,13–15^. Yet, the metabolic dimension of this cellular heterogeneity remains far less understood, even though metabolic reprogramming is a hallmark of GBM growth and treatment resistance.

Metabolic adaptations in GBM involve shifts in glycolysis and oxidative phosphorylation, changes in amino acid and nucleotide biosynthesis, and extensive rewiring of lipid metabolism^16–18^. Bulk metabolomics studies have demonstrated substantial metabolic variability among GBM tumors, including subtype-specific dependencies such as elevated purine synthesis in PN-like samples or lipid metabolic alterations in MES-like tumors^4,19^. However, bulk metabolomic profiles obscure the contributions of individual cell types that coexist within the tumor mass, including malignant states, immune cells, stromal populations, and vascular-associated cells. As a result, the mechanisms by which specific cell populations reprogram metabolism to support proliferation, invasion, immune evasion, or therapeutic resistance remain poorly defined.

Since current single-cell metabolomics technologies are technically constrained by low throughput, limited metabolite coverage, and the inability to amplify metabolites (and thus low sensitivity), alternative strategies are needed to infer metabolic states at cellular resolution^20,21^. One powerful avenue is the integration of scRNA-seq data with prior biochemical knowledge, allowing computational reconstruction of pathway activity, metabolic regulation, and flux distributions. Pathway-level inference approaches, gene regulatory network (GRN) analysis, and constraint-based modeling with genome-scale metabolic models (GEMs) each provide complementary windows onto cellular metabolism derived from transcriptomic readouts^20^. As these methods do not directly measure metabolites or reaction fluxes, their accuracy must be evaluated carefully, yet they offer a unique opportunity to map metabolic heterogeneity across thousands of individual cells.

Recent work by Wang et al. (2021) demonstrated how multiomic integration across 99 treatment-naive GBM samples can identify metabolic features associated with PN-like, CL-like, and MES-like GBMs, revealing subtype-specific vulnerabilities^4^. Building on these insights, computational approaches applied to single-cell expression data may enable the deconvolution of metabolic programs at the level of specific malignant states and non-malignant TME compartments, capturing heterogeneity that bulk profiling cannot resolve.

In this study, we investigate metabolic dysregulation across GBM cell populations using single-cell transcriptomic data coupled with complementary computational frameworks. We compared three methodological pillars. First, metabolic pathway activity analysis is used to identify pathways whose component genes exhibit coordinated up-or downregulation across malignant and non-malignant cell types, enabling assessment of intra- and inter-tumoral metabolic diversity^22^. Second, gene regulatory network inference using the SCENIC framework is applied to reconstruct transcription factor-driven regulatory programs controlling metabolic enzymes^23^. This approach highlights cell-state-specific regulators that may drive metabolic shifts within the tumor ecosystem. Third, we use a state-of-the-art metabolic flux prediction tool, scFEA, to estimate reaction activity and flux potentials at single-cell resolution based on genome-scale metabolic models^24^. These methods aim to infer the metabolic capacities of distinct GBM cell states, offering predictions on which metabolic pathways are likely to be active in individual cells.

Together, these three approaches provide complementary perspectives on GBM metabolism: pathway-level trends, regulatory drivers, and predicted reaction fluxes. By applying them systematically to the same scRNA-seq datasets, our study provides a comprehensive view of metabolic heterogeneity within GBM at single-cell resolution. This strategy contributes to understanding how malignant and non-malignant cell types reprogram metabolism in distinct ways and highlights metabolic dependencies that may offer opportunities for therapeutic intervention. As computational approaches for single-cell metabolic inference continue to evolve, systematic cross-validation against experimental metabolomics data will be essential to refine predictive accuracy and enhance the translational relevance of single-cell multiomics analysis in GBM and other heterogeneous cancers.

## Results

### Exploration of inter- and intra-patient glioblastoma tumor heterogeneity

For this study we explored snRNA-seq data of a set of individual GBM patients from the publicly available Wang dataset (2021). In the original study, the authors performed an integrated analysis of genomic, proteomic, post-translational modifications, and metabolomic data on 99 treatment-naive GBM samples^4^. Based on these bulk-level multiomics measurements, they clustered the GBM IDH-WT samples into 3 multiomic subtypes: nmf1 (PN-like), nmf 2 (MES-like), and nmf 3 (CL-like). Biologically, nmf1 was enriched for synaptic vesicle cycle, neurotransmitter transport and amino acid metabolism; nmf2 was enriched for innate immune response and glycolysis; and nmf3 was enriched for mRNA splicing, RNA metabolism, DNA repair and histone acetylation^4^. For a subset of these samples (18), the authors additionally provided snRNA-seq data (Suppl. Fig. 1).

We performed independent preprocessing and quality control on each sample. To ensure robust downstream metabolic inference and reliable cross-sample comparisons, we selected a subset of 6 high-quality samples (2 per Wang multiomic subtype). Samples were included based on: 1) sufficient cell numbers after QC filtering, 2) recovery of canonical cell-type marker genes described in the original study, and 3) consistent representation of major malignant and microenvironment cell populations in distinguishable cell clusters. In order to maximally retain inter-patient tumor heterogeneity, we analyzed each sample individually without an integration step.

Consistent with the observations of the original Wang study, tumor-associated macrophages (TAMs) represented the predominant non-neoplastic cell population in the GBM tumor-microenvironment (TME) across all 6 samples, typically followed in abundance by oligodendrocytes or their progenitors. A small population of monocytes was detected in 4 samples based on mapping to the GBMap reference atlas^25^, although these cells largely co-embedded within the TAM cluster, suggesting transcriptional similarity or transitional states (Fig. 1).

**Figure 1.**
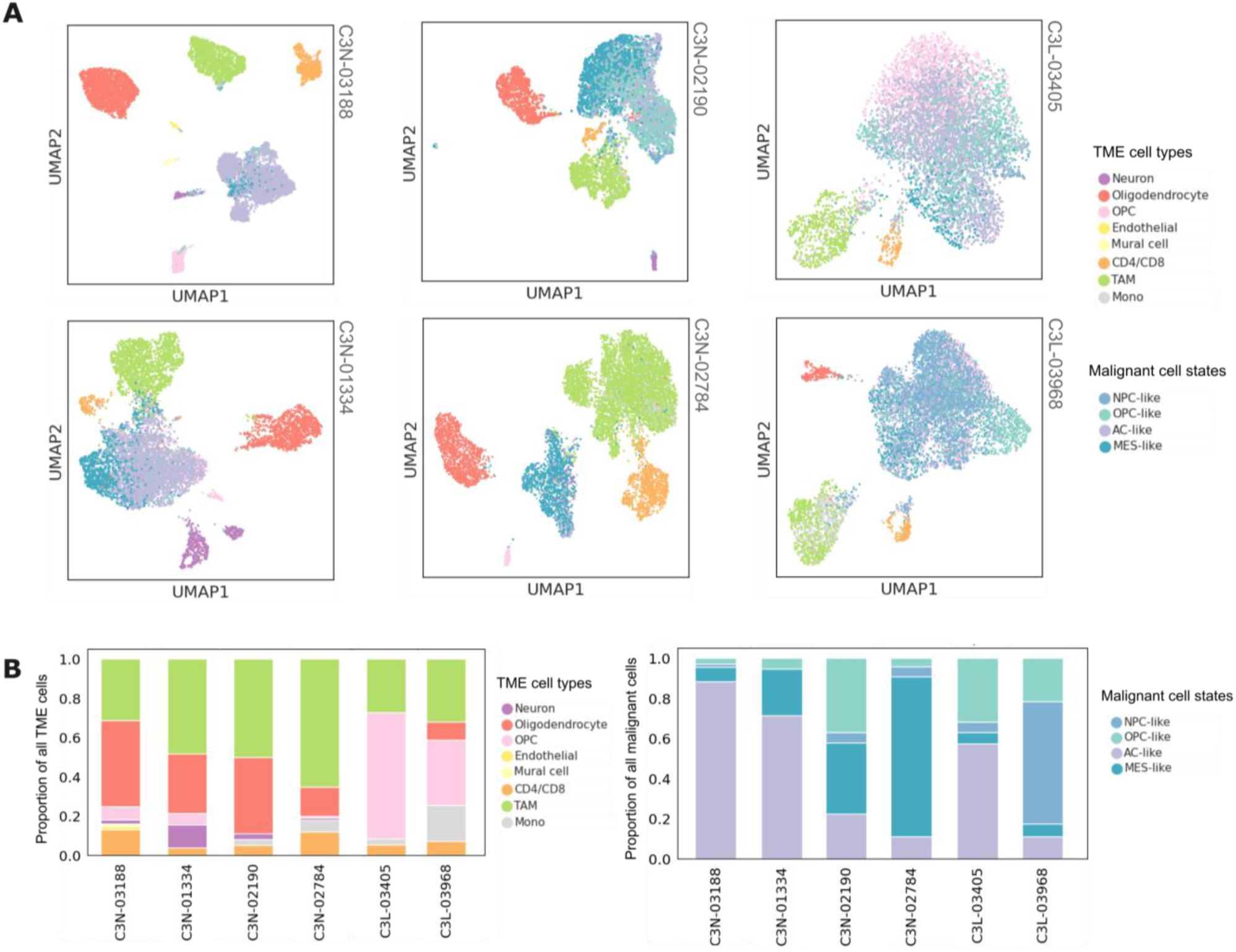
Inter- and intra-patient glioblastoma tumor heterogeneity. a) Uniform manifold approximation and projection (UMAP) for dimension reduction based on gene expression of cells in each of the 6 GBM samples colored according to the GBmap cell type/malignant cell state annotation. Sample IDs are provided on the top right of each respective UMAP. b) Proportions of each TME cell type in the TME compartment and each malignant cell state in the malignant compartment for each GBM sample. (OPC: oligodendrocyte progenitor cell; TAM: tumor-associated macrophage; Mono: monocyte; NPC: neural progenitor cell; AC: astrocyte; MES: mesenchymal)

Despite these shared broad patterns across all samples, GBM tissue composition remains highly diverse. And depending on patient background, sampling location, and disease stage, substantial variation occurs in both the presence and relative abundance of cellular populations across samples. This variability was particularly evident in the malignant cell compartment. Malignant cells were annotated based on the four Neftel cell states: NPC-like, OPC-like, AC-like, and MES-like. Samples belonging to the nmf1 multiomic subtype are dominated by AC-like malignant cell states, whereas nmf2 samples are characterized by a strong enrichment of MES-like cells. In contrast, samples of the nmf 3 subtype exhibit a more diverse mixture of malignant cell states but show highly similar non-malignant cellular compositions, including a prominent oligodendrocyte progenitor cell (OPC) population (Figure 1B). Together, these patterns highlight the interplay between shared TME features and subtype-specific malignant cell-state landscapes across samples. Such a pronounced cell type and malignant cell state heterogeneity implies that the metabolic wiring of GBM cannot be captured by bulk measurements or by pooling together single-cell data across patients. Metabolic pathway usage, regulatory programs, and flux distributions are likely to differ not only between cell types, but also between patients and tumor subtypes. This motivated a sample-specific single-cell metabolic modeling strategy, in which each patient’s dataset is analyzed independently to preserve the biological variation inherent to their cellular composition and malignant state architecture. In what follows, we applied and directly compared 3 conceptually distinct methods on all samples: pathway activity scoring, gene regulatory network inference, and metabolic flux prediction. To enable fair and uniform comparison, all methods were mapped onto a shared pathway annotation framework from the Human1 GEM^26^.

### Metabolic pathway activities reveal cell-state-specific activity patterns

To characterize the metabolic landscape across different cell populations of the selected GBM samples, we first assessed pathway-level metabolic activity across malignant and non-malignant cell populations. We quantified metabolic pathway activity using the approach of Locasale^22^ in combination with curated metabolic pathway gene sets (genes encoding for metabolic enzymes) derived from the Human1 GEM model. The Human1 pathways were manually evaluated, consolidated to reduce redundancy, filtered for sufficient gene coverage, and scored per cell type using normalized expression values with gene weighting to account for multi-pathway genes (Methods). This approach produced robust pathway activity estimates for each sample, enabling comparison of metabolic programs across the major cellular populations identified.

When ranking pathways by their maximum activity across cell types within each sample and comparing these sample-wise rankings, a recurrent set of highly active pathways was observed (Fig. 2). Protein degradation, drug metabolism, protein assembly, and even-chain fatty acid desaturation consistently appeared among the top-ranking pathways across samples, being present in the top 5 pathways in at least half of the samples. This indicates a strong and conserved metabolic engagement in these processes across patients. When examining cell type specificity of these top-ranked pathways, we observed clear patterns: protein degradation/assembly and drug metabolism were predominantly active in TAMs and monocytes, whereas even-chain fatty acid desaturation was mainly active in oligodendrocytes or their progenitors (Fig. 2c).

**Figure 2.**
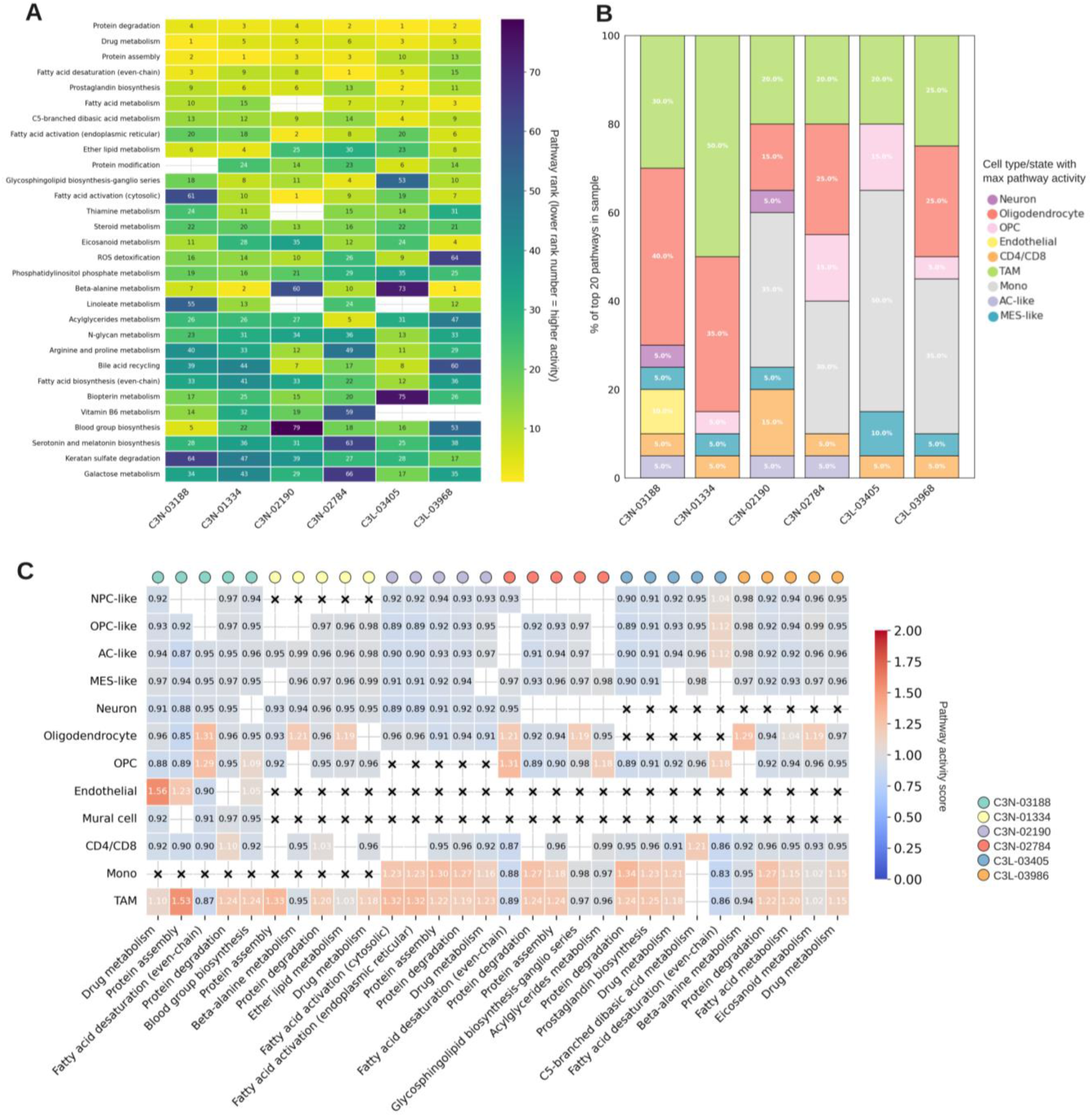
Metabolic pathway activities across samples and cell types/states. a) Heatmap displaying pathway rankings across samples (x-axis). Within each sample, pathways were ranked based on their maximum activity across cell types/states. These sample-specific ranks were subsequently aggregated to identify the top 30 most consistently highly ranked pathways across all samples. Lower rank number indicates higher pathway activity. b) Per sample, each pathway has a maximum activity in a particular cell type/state. Stacked bar plots show for each sample the proportion o the top 20 pathways whose highest activity was attributed to each cell type/state, colored accordingly. c) Pathway activity scores for the top 5 pathways in each of the 6 samples. Empty entries refers to non-significant activity values for a given pathway-cell type/state combination. A black cross indicates the absence of that cell type/state (y-axis) in the respective sample. Samples are distinguished by colored dots along the top of the heatmap.

Even when expanding the ranking window to the top 20 pathways per sample, we further observed the influential role of TAMs, monocytes, and oligodendrocytes in driving high pathway activity scores (Fig. 2b). Within this broader set, additional fatty acid- and lipid-related metabolic pathways became apparent, including prostaglandin biosynthesis (OPC-active), fatty acid metabolism (TAM- and monocyte-active), fatty acid activation (TAM- and monocyte-active), ether lipid metabolism (oligodendrocyte-active), glycosphingolipid biosynthesis of ganglio-series (oligodendrocyte-active), as well as steroid and eicosanoid metabolism (oligodendrocyte-active).

In contrast, most high-ranking pathways showed only limited activity in any of the malignant cell states. Only two pathways were still moderately ranked across samples (appearing in the top 20 of at least half the samples) and displayed their highest activity in MES-like malignant cells, namely protein modification and ROS detoxification.

Finally, a small number of pathways appeared among the top 10 in only one or two samples such as beta-alanine metabolism, acylglyceride metabolism, and blood group biosynthesis, suggesting rather sample-specific metabolic features.

As reported by Wang et al., samples from the nmf2 (MES-like) multiome subtype were characterized by an enrichment in innate immune response and glycolysis. Examining these same samples in our study (C3N-02190 and C3N-02784), we found that a large proportion of the top 20 most active pathways in each sample were driven predominantly by TAMs and monocytes, suggesting that this innate immune enrichment is accompanied by pronounced metabolic activity in these cell populations (Fig. 2b). Zooming in more specifically on the TAM compartment within both nmf2 samples, fatty acid-related pathways emerged as the most highly active, with fatty acid activation and fatty acid metabolism ranking among the top pathways (Suppl. Fig. 2). This points to a metabolically distinct TAM phenotype in MES-like tumors, where lipid metabolism may play a particularly prominent.

### Regulatory network inference highlights TF-driven control of a set of metabolic programs

While pathway-level activity profiling provided a first view of the intra- and intertumoral metabolic heterogeneity across samples, it did not yet explain how these metabolic programs are mechanistically regulated. To investigate the potential gene regulatory mechanisms that may drive these metabolic states, we next inferred transcription factor (TF)-centered regulatory networks using SCENIC^23^. SCENIC reconstructs regulons, i.e. TFs and their predicted target genes, and quantifies their activity at single-cell resolution. We used these regulon activity profiles to identify the dominant regulatory programs in each sample and to assess their cell type specificity. We then evaluated whether cell type-specific regulons were enriched for metabolic genes, enabling us to link regulatory programs to metabolic phenotypes.

Running SCENIC independently on each patient sample yielded between 41 and 65 regulons per sample (Suppl. Fig. 3b). To compare regulon content across samples, regulons were matched solely based on their TF name, since SCENIC reconstructs patient-specific target gene sets and thus regulons with the same TF label are not guaranteed to share an identical target gene composition. Only a minority of regulons were detected across all patients (RUNX3 and PRDM1), with the majority of regulons (54,3%) being specific to a single sample (Suppl. Fig. 3a-c). The MES-like subtype sample C3N-02190 comprised the largest set of sample-specific regulons (Suppl. Fig. 3b), with most of them showing the highest regulon activity in the MES-like malignant cell state (TCF7L2, RFX2, NR2F6, KLF5, RELB). This suggests substantial inter-patient regulatory heterogeneity and again reinforces the need for a per-sample analysis.

Despite the variability in regulon composition, regulon activity patterns also revealed clear and reproducible cell type specific signatures across some samples. When looking at regulons found in at least half of the samples, TAMs and monocytes showed high activity of regulators involved in myeloid lineage identity and inflammatory signaling (SPI1, IRF8, IRF5, RUNX1, CEBPB, CEBPD, ETS2, FLI1, NFATC2, ATF3), hypoxia (HIF1A), environmental stress sensing and immediate-early responses (FOS, FOSB, FOSL2, JUNB, ATF3), and early-response and chromatin modeling (EGR2, EGR3, ATF3, PRDM1). Oligodendrocytes were characterized by strong activity of the FOXO1 regulon. And the malignant cells showed strong activity of regulons involved in proliferation (E2F2 and E2F7), stem-state maintenance (SOX4, TCF7L1), and endothelial plasticity (ETV2). These malignant cell-specific regulons were each time most highly active in different malignant cell states when comparing across different samples (Suppl. Fig. 3d). Detailed regulon activity heatmaps per sample are shown in Suppl. Fig. 4.

To connect the regulatory programs to metabolic phenotypes, we quantified the overlap between each regulon’s target genes and the curated metabolic pathway gene sets of Human1 as used in our first approach. Here we essentially construct tripartite networks (three different node types) connecting TFs to target genes to metabolic pathways. Each sample yielded a regulon × pathway overlap matrix, where each entry represents the number of target genes of a regulon found in a specific pathway. The larger this number, the more a specific TF controls the expression of that pathway’s enzymes.

Having established which regulons connect to which metabolic pathways and how many genes within a single pathway can be regulated by one or multiple TFs we first identified global patterns of this metabolic regulation across all samples. We wanted to identify regulons with similar metabolic pathway regulation patterns. Such regulons may participate in shared biological processes or coordinated metabolic programs.

To assess this similarity, we adapted the principle of the connection specificity index (CSI). Originally, the CSI is used in GRN contexts with TF-target gene interactions as a context-dependent graph metric that mitigates the effect of nonspecific interactions by assessing the similarity between two regulons based on their target gene interaction profiles across the entire network^27^. We redefined this CSI concept for our metabolic analysis by transforming tripartite networks into weighted bipartite networks. Specifically, we drew a directed edge between each TF and all metabolic pathways with which it was connected through target genes, with edge weights corresponding to the number of overlapping target genes. In this reformulated framework, the CSI score between a pair of regulons now reflects the degree to which both regulons are involved in regulating similar metabolic pathways (Fig. 4a). Operationally, this was achieved by using the regulon × pathway matrix described above, where each entry represents the number of overlapping genes, and then calculating CSI correlation values between all regulon pairs.

This CSI scoring gave rise to a set of regulon modules which seemed to have a very similar pathway regulation profile. For example, we identified a module of regulons for which ASCL1, LHX9, EVX2, and ONECUT2 were specifically found in only one sample (MES-like sample C3N-02190). Within this same module we also found TBX21 (present in all but sample C3N-02784 and C3L-03968) and TAL1 (found in C3N-02190 and C3L-03968). Within this module the regulons were found to be mainly highly active in monocytes and TAMs. Importantly, not all regulon modules with high CSI scores were found to be sample-specific. We additionally identified a considerable module existing of 30 different regulons which showed activity across nearly all samples. Strikingly, despite the pronounced inter-sample heterogeneity in cellular composition and malignant cell states described above, these 30 regulons point to a conserved transcriptional core spanning both immune and malignant compartments. In myeloid cells this is centered on inflammatory programs: NFκB, AP-1, and IRF-driven inflammatory activation in TAMs and monocytes is strongly linked to a pro-inflammatory metabolic rewiring specifically enhanced glycolysis (Warburg-like metabolism) and itaconate/succinate accumulation, which are hallmarks of M1-like macrophage polarization^28^. CEBPB is also well-known regulator of lipid metabolism in myeloid cells^29^, consistent with the fatty acid pathway activity already described in the previous TAM compartment of nmf2 samples. In malignant cells on the other hand the transcriptional core is centered on hypoxia- and proliferation-driven programs: HIF1A as a master regulator of metabolic adaptation to hypoxia directly drives glycolysis, suppresses oxidative phosphorylation, and promotes angiogenesis. The E2F factors and MYBL1 are tied to nucleotide synthesis and one-carbon metabolism to fuel rapid proliferation.

To more specifically illustrate how regulons connect to metabolic pathways, we focused on a set of regulons specifically active in TAMs of the two MES-like samples (C3N-02190 and C3N-02784), i.e. CEBPD, FOSL2, HIF1A, IRF5, and PRDM1 (Fig. 3a). For both samples, we identified a recurring set of pathways for which a substantial number of enzymatic genes were inferred to be regulated by several or all of these TFs. When examining the broader metabolic categories to which these pathways belong, we found an enrichment of lipid and fatty acid metabolism-related pathways (Fig. 3b). For example, in sample C3N-02784 all five TFs were predicted to regulate a significant number of enzymatic genes within the sphingolipid and inositol phosphate metabolic pathways. More broadly, the pathway overlap matrix revealed that PRDM1 consistently shows the largest absolute overlaps across nearly all pathways presented. In this sample all five TFs showed substantial overlaps particularly in lipid-related pathways such as sphingolipid, glycerophospholipid, glycerolipid, and arachidonic acid metabolism, suggesting that these TFs converge on a shared lipid-regulatory program in TAMs (Fig. 3b). The network visualization further highlights this convergence, showing that the same set of lipid-related pathways are targeted by multiple TFs simultaneously. Moreover, these TFs also seem to regulate each other, pointing to a coordinately regulated lipid metabolic program (Fig. 3c).

**Figure 3.**
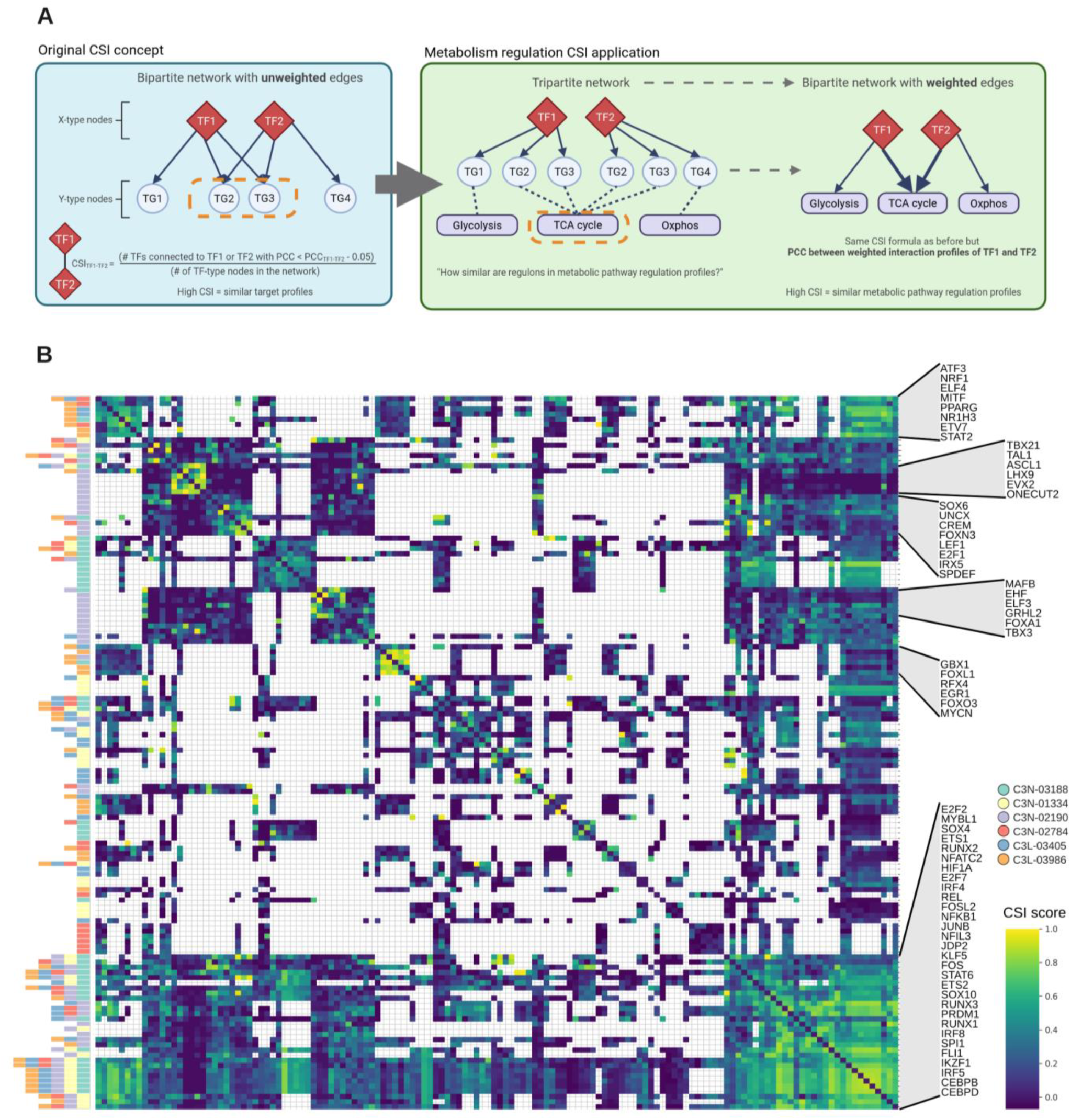
CSI-based quantification of metabolic regulatory similarity across regulons and samples. a) Schematic illustration of the CSI computation adapted for metabolic pathway regulation. b) Aggregated CSI matrix displaying pairwise regulon similarity scores across all samples. Each entry represents the CSI score between a pair of regulons with higher values indicating more similar metabolic pathway regulation profiles. Regulons are ordered by hierarchical clustering to reveal modules of co-regulating TFs. The TFs comprising different modules are annotated on the right. Annotation bars on the left show in which and how many samples the regulon was inferred. Empty entries indicate regulon pairs that were not co-observed in any of the samples and therefore received no CSI score.

To eliminate the effect of regulon size on these absolute overlap values, we additionally calculated overlap proportions, defined as the fraction of a regulon’s total target genes that also encode enzymes in a specific pathway. The consensus of overlap proportions across all samples can be consulted in Suppl. Fig. 6.

### Single-cell flux estimation identifies sub-pathway level metabolic dependencies in glioblastoma

While the previous two approaches were informative, their analyses evaluate pathways in isolation, since we merely used the Human1 GEM model as a collection of metabolic pathway gene sets and thereby we disregarded the highly complex and interconnected nature of the complete metabolic network. Essentially, we did not fully exploit the network characteristics of this metabolic model. In our final approach, this limitation was addressed by computing metabolic reaction fluxes.

To accomplish this we used the scFEA flux prediction tool^24^. This tool leverages an adapted form of flux balance analysis in convergence with prior knowledge in the form of a reduced GEM to predict cell-wise metabolic flux distributions by integrating single cell transcriptomics data with GEMs. scFEA predicts flux for each of the 168 metabolic modules across individual cells, with modules defined as groups of connected metabolic reactions ensuring limited connections of intermediate metabolites to other modules. Supermodules in the scFEA results refer to manually curated groups of modules with similar functions, as specified in the scFEA paper^24^. These supermodules facilitate the linkage of individual reactions or reaction modules to functional pathway annotations.

We first visualized the similarity of single cells based on their flux prediction profiles in UMAPs for each of the 6 samples. Essentially, each cell was positioned based on the similarity of its overall metabolic profile to other cells. Across all samples, the predominant cell populations of the TME grouped into distinct clusters according to their GBmap cell type annotation, particularly evident for TAMs and oligodendrocytes (Fig 5A). This suggests that different TME cell populations employ distinct metabolic programs. In contrast, malignant cells did not clearly separate into distinct clusters on the UMAPs according to their expression-based Neftel cell states.

**Figure 4.**
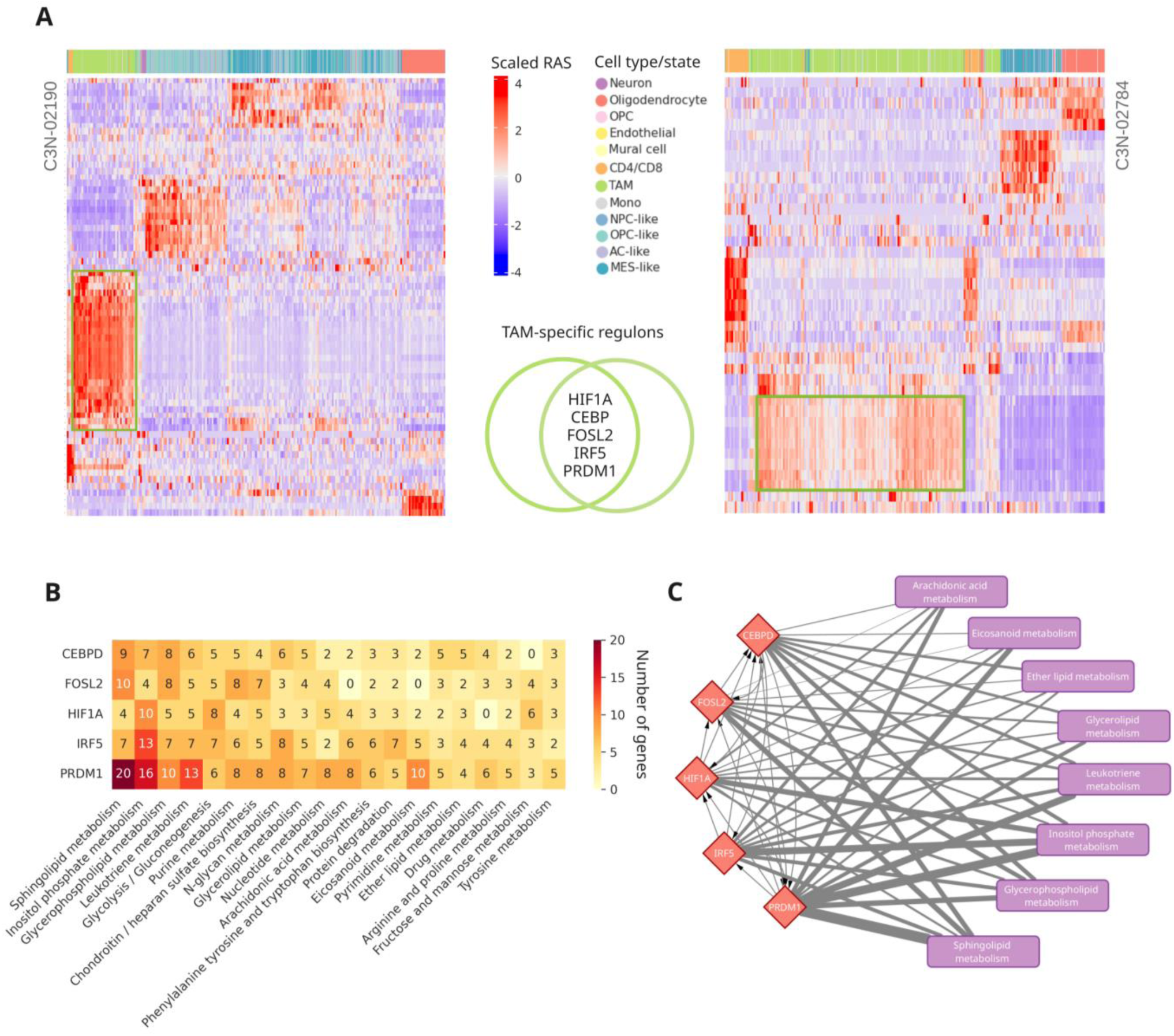
TAM-specific regulon-to-pathway connectivity in MES-like samples. a) Scaled regulon activity scores (RAS) across all cells for the 2 nmf2 (MES-like) samples C3N-02190 and C3N-02784. Cells are colored by cell type/state along the top color bar. Rows and columns of the heatmap are hierarchically clustered. In between heatmaps are the 5 TAM-specific regulons recurrently identified across both samples. b) Heatmap displaying the absolute number of target genes per regulon (rows) that overlap with enzyme-coding genes of each Human1 metabolic pathway (columns) for sample C3N-02784. c) Network visualization of the regulon-pathway connectivity for the regulons shown in a) and b) for sample C3N-02784. Red nodes represent TFs, purple nodes represent metabolic pathways, and edge width represents number of overlapping target genes between regulon and pathway.

**Figure 5.**
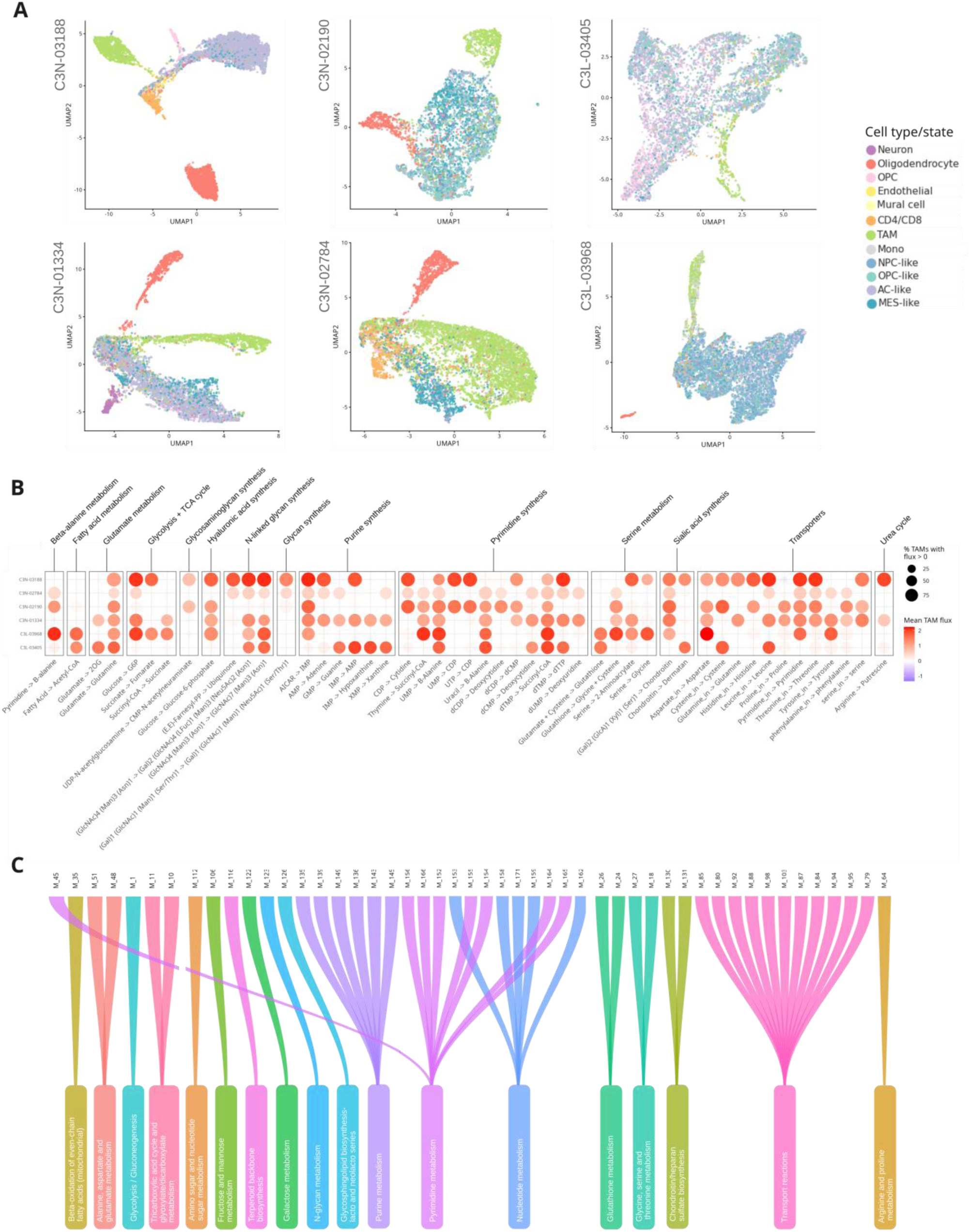
Single-cell metabolic flux predictions across TME cell populations and TAM-specific reaction modules. a) UMAP embeddings of single cells flux predictions from each of the 6 samples. Cells are positioned according to similarity of their predicted metabolic flux profiles across 168 scFEA reaction modules. Cells are colored by GBmap cell type/state annotation. b) Bubble plot summarizing TAM-specific reaction modules across all samples. Each row represents a sample and each column represents an scFEA reaction module, grouped by supermodule annotation (indicated on top). Bubble color reflects mean TAM flux of all TAM cells in the respective sample and through the respective reaction module. Bubble size indicates the % of TAM cells with positive flux in that sample. Only modules meeting TAM-specificity criteria in at least 2 samples are shown (Cohen’s d > 0.8 and >50% of TAMs with positive flux). c) Mapping between TAM-specific scFEA reaction modules (top) and Human1 GEM pathway annotation (bottom). Each ribbon connects an scFEA module from b) to its corresponding Human1 pathway.

Since the TAM population formed a distinct cluster across all samples based on predicted flux profiles, we further examined which individual reaction modules showed specifically high flux through TAM cells and whether these were enriched for specific supermodules. For each sample, we extracted the scFEA reaction modules that showed specifically high flux through TAMs compared to other cell types and malignant states. Across the samples, we observed high flux through a large set of purine and pyrimidine synthesis reaction modules, indicating strong involvement in nucleotide biosynthesis. Within the purine synthesis supermodule, the AICAR -> IMP module showed high TAM-specific flux across almost all samples. In the pyrimidine synthesis supermodule, the UMP -> B-alanine and dTMP -> succinyl-CoA modules exhibited TAM-specific high fluxes in all studied samples, suggesting global importance. We additionally identified strong flux through many influx transport reactions, suggesting that TAMs are not only metabolically active internally, but also strongly interact with their environment. Another highly relevant metabolic reaction module found across all samples was the conversion of glutamate to glutamine. In GBM, TAMs are notorious to use glutamine to fuel their pro-tumorigenic, immunosuppressive state by converting glutamate to glutamine, creating a nutrient sink that starves T cells^30^.

Other TAM-specific high reaction fluxes were rather sample-specific or present only in a subset of samples. For instance, the Fatty Acid -> Acetyl-CoA module was found to be TAM-specific only in the two samples of the nmf3 subtype (CL-like). Surprisingly, no fatty acid related metabolic modules were identified in the two nmf2 MES-like samples. Another module specifically found in the two nmf3 samples was Glutamate + Cysteine -> Glutathione. Glutathione is known to be important for malignant cell survival, acting as an antioxidant that reacts with most reactive oxygen species^31^. Additionally, one glycan synthesis reaction module was found specifically in the two nmf1 (PN-like) samples.

The challenge of interpreting and comparing these scFEA results to the previous two approaches is that the supermodule annotations often deviate significantly from the previously used Human1 GEM pathway annotations (e.g. supermodule “Glycolysis+TCA cycle” vs Human1 pathway “Glycolysis/Gluconeogenesis”). Therefore, we mapped the individual reaction modules of scFEA to Human1 pathway annotations based on the enzymes catalyzing the reactions of each module (Suppl. Fig. 7). Applying this mapping to the TAM-specific reaction modules discussed above revealed that some reaction modules belonging to one supermodule also mapped to a single Human1 pathway (e.g. Transport reactions). Conversely, some reaction modules were grouped into one supermodule despite actually belonging to multiple different Human1 pathways (e.g. scFEA “Serine metabolism” supermodule vs Human1 “Glutathione metabolism” and “Glycine, serine and threonine metabolism”)(Fig. 5c).

### Cross-method comparison

The three analytical strategies applied in this study provide complementary perspectives on cellular metabolism, each operating at a different conceptual layer of resolution. The metabolic pathway activity scores offer a broad overview of transcriptionally driven pathway engagement at the cell type level. This approach is well suited for identifying which major metabolic pathways are most active across the cellular landscape of each sample, and it robustly highlights consistently active processes, particularly within TAMs, monocytes, and oligodendrocyte-lineage cells. However, it does not operate at single-cell resolution and does not incorporate network structure, limiting its ability to capture regulatory or reaction-level specificity.

The SCENIC-based metabolic regulation analysis extends beyond pathway abundance by identifying transcription factors whose regulons contain metabolic enzyme genes. Because regulon activity is quantified per single cell, this method reveals transcriptional programs underlying metabolic differences and identifies groups of regulons with similar metabolic targets. While regulon composition varies across samples, regulon activity maps and CSI-derived modules help uncover both sample-specific and recurrent regulatory patterns, providing mechanistic context that is not accessible from pathway scores alone.

In contrast, scFEA flux prediction focuses on reaction activity rather than gene or regulon activity, estimating feasible metabolic flux distributions for each cell based on a reduced genome-scale model. This method captures network constraints and sub-pathway structure, enabling the identification of cell type–specific flux patterns and metabolic dependencies that may not be apparent from transcript-level data alone. Because scFEA uses its own modular reaction framework, an additional mapping step is required to align its outputs with Human1 pathway annotations used in the other analyses.

Together, these methods span a hierarchy that provides increasing granularity and mechanistic insight: Metabolic pathways → regulatory programs controlling enzymes in the pathway → predicted reaction fluxes.

## Discussion

Glioblastoma remains one of the most adaptive and treatment-resistant human cancers. Its cancer cells undergo extensive metabolic reprogramming to meet the increased demands of rapid growth and to survive in an often oxygen- and nutrient-deficient TME. In this study, we systematically compared 3 transcriptomics-based approaches to interrogate metabolic diversity across malignant and non-malignant cell populations in GBM, namely metabolic pathway activity scoring, TF-based metabolic regulation analysis, and single-cell flux prediction. Together these analyses reveal both converging and distinct aspects of tumor metabolism, highlighting the need for multi-layered frameworks to fully interpret metabolic rewiring in GBM.

A central and consistent finding across all 3 analytical layers is the metabolic prominence of TAMs within the GBM TME. This myeloid population drove the highest pathway activity scores, showed the most coherent regulon activity modules, and formed the most clearly distinct cluster in flux-based UMAP space. This convergence strongly supports the view that TAMs are not merely bystanders, but metabolically dominant players in GBM. Pro-tumoral TAMs are better adapted to the nutrient-depleted TME (low glucose) as they rely on fatty acid oxidation for energy production to enable a sustained immunosuppressive function^32^. Our findings are consistent with this picture and add mechanistic resolution by identifying specific pathways and regulatory programs driving this phenotype at single-cell resolution.

Notably, our analyses identified the prominent engagement of lipid and fatty acid metabolism pathways in TAMs, especially in MES-like tumors. Pathway activity scoring identified fatty acid activation and fatty acid metabolism among the top TAM-specific pathways in nmf2 samples, and the GRN analysis revealed that 5 TAM-specific TFs (CEBPD, FOSL2, HIF1A, IRF5, and PRDM1) converge on a shared lipid-regulatory program with sphingolipid, glycerophospholipid, and arachidonic acid metabolism being coordinately targeted. This finding is consistent with recent work showing that lipid-laden macrophages phagocytose myelin debris through the lipid receptor CD36, shifting from an inflammatory to an immunosuppressive phenotype. In this way they reciprocally recycle lipids to fuel GBM progression and this interaction is particularly prevalent in MES-like tumors^33^. The broader importance of CEBP-family activity in orchestrating the pro-tumorigenic myeloid niche in GBM is further supported by recent work showing that CEBPB expression in GBM tumor cells drives M2 TAM recruitment through SPP1-mediated signaling and correlates with poor prognosis^34^.

The GRN analysis also identified a conserved transcriptional core of 30 regulons spanning both immune and malignant compartments across nearly all samples. This module of regulons was centered on inflammatory programs (NFκB, AP-1, IRF family) in myeloid cells and hypoxia- and proliferation-driven programs in malignant cells. We recovered HIF1A as an active regulon in TAMs across multiple samples. This is a well established master regulator of the hypoxic response in GBM, promoting malignant progression by inhibiting apoptosis, increasing proliferation, and driving stemness under hypoxic conditions^35^. Importantly, we identified the E2F family members as conserved regulons in the malignant compartment. They are known to drive nucleotide biosynthesis programs. Promoters of nucleotide biosynthesis genes share common binding sites for TFs including E2F factors which have been experimentally verified as regulators of this program^36^. This finding connects directly to the flux analysis results, in which purine and pyrimidine synthesis modules showed the strongest and most consistent TAM-specific flux signals across all samples. However, elevated nucleotide biosynthesis has also been implicated in GBM malignant cells themselves, where high rates of de novo purine synthesis promote the maintenance and tumorigenic capacity of glioma-initiating cells and contribute to therapy resistance and tumor recurrence^37^.

The flux analysis additionally identified strong glutamate-to-glutamine conversion as a recurrent TAM-specific flux across all samples. This aligns with recent literature describing a metabolic symbiosis between TAMs and GBM cells in which, in response to a glutamate-rich TME, TAMs upregulate glutaminase activity and synthesize glutamine from glutamate, effectively supplying GBM cells with a nutrient they are critically dependent on. GBM cells in turn exhibit aggressive glutamine uptake that depletes this nutrient from the TME, potentially impairing CD8+ T cell function and contributing to immune evasion. This dynamic has been described as a glutamine-driven "metabolic tug-of-war" between tumor and immune cells^38^.

Fatty acid oxidation (fatty acid → acetyl-CoA) on the other hand emerged as TAM-specific exclusively in nmf3 (CL-like) samples, whereas no fatty acid flux modules were identified in nmf2 MES-like TAMs despite strong fatty acid pathway activity and regulon signals in those same samples. This apparent discrepancy likely reflects a methodological difference in sensitivity: the pathway activity and GRN approach capture transcriptional engagement across all enzyme-coding genes in a pathway, whereas scFEA flux prediction captures only linearizable reaction modules meeting strict stoichiometric constraints. This suggests the fatty acid metabolic program in MES-like TAMs may be distributed across reaction steps that do not form a linearizable module in the scFEA framework, underscoring the complementary nature of the 3 approaches.

An important implication of our findings is that metabolic reprogramming in GBM cannot be accurately captured using a single transcriptomics-derived method. Each analytical layer provides a distinct vantage point: pathway activity summarizes upregulated metabolic themes, regulon analysis exposes the transcriptional logic underlying these themes, and flux modeling incorporates biochemical constraints to approximate feasible metabolic behavior on (near) reaction level. The areas where these methods converge are likely to represent robust features of GBM metabolism, particularly the metabolic prominence of TAMs. In contrast, areas of divergence highlight biological complexity or methodological limitations that warrant further investigation.

Practically, this integrated strategy has implication for therapeutic development. Targeting lipid metabolism in TAMs represents a promising strategy to modulate the immunosuppressive TME and enhance response to standard therapies^39^. Our identification of specific TFs as coordinated drivers of this lipid program in MES-like TAMs provides a regulatory entry point for such strategies. Similarly, the consistent TAM-specific purine and pyrimidine synthesis flux signals suggest that nucleotide biosynthesis inhibitors which have already been explored in the context of GBM malignant cells, might also modulate TAM function within the TME. As computational approaches for single-cell metabolic inference continue to mature, systematic cross-validation against spatial or single-cell metabolomics will be essential to refine the predictive accuracy and translational relevance of these integrative frameworks in GBM and potentially other heterogeneous cancers.

## Materials & methods

### Data retrieval

Single-nucleus RNA-seq (snRNA-seq) was obtained from Wang et al. (2021), who performed multi-omic profiling of treatment- naïve GBMs collected by the Clinical Proteomic Tumor Analysis Consortium (CPTAC)^4^. From this cohort, 18 IDH-wildtype adult GBM samples (9 male, 9 female; age range 24–77) were analyzed. Based on multiomic NMF clustering, Wang et al. classified samples into 3 subtypes: nmf1 (proneural-like), nmf2 (mesenchymal-like), and nmf3 (classical-like), broadly concordant with TCGA expression subtypes though with some reassignments. A subset of samples exhibited mixed subtype identity based on low NMF membership scores (Suppl. Fig. 1).

snRNA-seq was performed by Wang et al. on the 10X Chromium platform (∼20,000 nuclei per sample), with FASTQ files processed using Cell Ranger v3.1.0 against a customized pre-mRNA GRCh38 genome reference. Unfiltered feature-barcode matrices (count matrix) were used as input for further downstream analysis.

### snRNA-seq preprocessing and quality control

Count matrices were preprocessed individually per sample using SCANPY^40^ (v1.9.8). Cells that fell into any one of the following categories were filtered out: cells with less than 200 genes expressed (possible debris); cells with less than 1000 total transcript counts; cells with more than 10,000 genes expressed; cells with more than 10,000 UMIs; or cells with a proportion of mitochondrial transcript counts over the total transcript counts higher than 10%. Genes were filtered if they were expressed in less than 3 cells. Filtered matrices were normalized by median count depth followed by log1p transformation. The 6,000 most highly variable genes (HVGs) were identified using the Seurat flavor^41^, followed by PCA (n=50 components, arpack solver), neighborhood graph construction (n_neighbors=20, n_pcs=30), UMAP embedding, and Leiden clustering (resolution=0.5) (Traag et al., 2018).

Six samples were selected for downstream analysis, 2 per multiomic subtype, based on manageable post-filtering cell counts (≤10,000), low mitochondrial gene fractions, well resolved UMAP clusters, and adequate coverage of Wang et al. marker genes.

### Cell type and malignant cell state annotation

Cell type annotation was performed by combining unsupervised manual annotation with reference mapping against GBmap^25^, a curated GBM atlas spanning 26 datasets, 240 patients, and >1.1 million cells. For manual annotation, known marker genes from Wang et al. were visualized on UMAP embeddings and ambiguous clusters were resolved using the top 5 differentially expressed genes queried against CellMarker 2.0^42^ and the Human Protein Atlas^43^.

For reference mapping, filtered count matrices were imported into R (v4.3.2) and processed as Seurat (v5.1.0) objects. Transfer anchors between query and GBmap reference were identified using FindTransferAnchors (50 PCs, log-normalized data), and cell type labels were transferred using TransferData. Query embeddings were projected onto the reference UMAP via IntegrateEmbeddings and ProjectUMAP, and mapping quality was assessed using MappingScore.

### Metabolic pathway activity analysis

Metabolic pathway gene sets were derived from the Human1 GEM. Gene-to-pathway mappings were constructed using information from the RXNS (reactions) and GENES tables, converting all Ensembl gene identifiers to gene symbols. To reduce redundancy among pathway definitions, pathways with identical gene sets were merged. In particular, all peroxisomal β-oxidation pathways, excluding the phytanic acid pathway, were consolidated into a single pathway designated “Beta oxidation of fatty acids (peroxisomal)”, and all mitochondrial β-oxidation pathways were merged into “Beta oxidation of fatty acids (mitochondrial)”. Likewise, redundant fatty-acid biosynthesis and elongation pathways were combined represented under “Fatty acid biosynthesis (unsaturated)”. Pathways represented by fewer than 3 genes in the scRNA-seq dataset were removed prior to scoring.

Because some genes participate in multiple pathways, a weighing scheme was implemented to prevent such genes from disproportionately influencing pathway scores. Each gene was assigned a weight inversely proportional to the number of pathways in which it appears. Pathway activity scores were computed for each cell type using normalized gene expression values. For each pathway, genes present in both the curated pathway set and the expression matrix were identified, and their expression values were integrated using the gene-specific weights. Cell type-specific mean expression values were then calculated to obtain weighted and unweighted pathway activity scores. To assess statistical significance, permutation testing was performed using 5000 random gene sets matched in size to each pathway, generating an empirical null distribution against which observed pathway scores were compared. This procedure yielded pathway-level activity estimates for each cell type along with empirical p-values reflecting the probability of observing the corresponding score under the null model.

### GRN inference and metabolic pathway regulation

For GRN inference SCENIC^23^ was run 10 times on each sample individually via the VSN Nextflow pipeline (https://github.com/vibsinglecell-nf/vsn-pipelines). Coexpression modules were inferred with GRNBoost2 using a list of manually curated human TFs of Lovering et al. (2021)^44^; regulon prediction with cisTarget, the motif and track databases outlined in the pbmc10k.vsn-pipelines.complete.config file (https://github.com/vib-singlecellnf/vsn-pipelines-examples/pbmc10k) were used. Other parameters for the cisTarget and the subsequent AUCell algorithm were the same as in the pbmc10k.vsn-pipelines.complete.config file. Per-sample consensus regulons were defined by retaining only TF-target edges present in ≥80% of runs. To subsequently study metabolic regulation, only regulon target genes encoding enzymes involved in Human1 metabolic pathways were considered.

To quantify the similarity of metabolic pathway regulation patterns between regulons, we adapted the Connection Specificity Index (CSI) as described by Bass et al.^27^. The CSI is a context-dependent graph metric that assesses the similarity between two nodes based on the specificity of their shared interaction partners, while mitigating the influence of nonspecific or promiscuous interactions. For each sample, we first constructed a weighted bipartite network from the regulon x pathway overlap matrix. In this network, regulons (TFs) and metabolic pathways constitute the two node types, with directed edges drawn from each regulon to all pathways for which at least one target gene overlap was identified. Edge weights correspond to the number of target genes in the regulon that encode enzymes in the respective pathway.

This CSI formulation assigns higher scores to regulon pairs that share connections to pathways that are themselves less promiscuously connected across the entire network, thereby emphasizing specific rather than generic metabolic regulation patterns. The resulting CSI values range from 0 (no similar pathway regulation) to 1 (identical pathway regulation patterns). For each sample, a regulon x regulon CSI matrix was computed, quantifying the pairwise similarity of metabolic regulatory strategies across all regulons in that sample.

### Metabolic flux prediction

To predict cell-wise flux the scFEA tool (v1.1.2) was applied to the scRNA-seq data of each sample individually. For large datasets, the authors recommend to only use gene expression data of the scFEA metabolic genes that will be used for flux computation. Expression matrices were subset to scFEA metabolic genes and run with default parameters (module_gene_m168.csv, cmMat_c70_m168.csv, sc_imputation=TRUE), yielding per-cell flux estimates across 168 metabolic modules.

To identify TAM-specific metabolic activity, per-module flux values were z-score normalized across cells within each sample. TAM specificity was quantified using Cohen’s d (TAM vs. non-TAM mean scaled flux divided by pooled SD), and the proportion of TAM cells with positive flux was computed. A module was classified as TAM-specific if Cohen’s d > 0.8 and > 50% of TAMs showed positive flux. Modules meeting these criteria in ≥2 samples were considered recurrently TAM-associated.

### Supermodule to Human1 pathway mapping

To enable direct comparison between scFEA flux estimates and the Human1-based pathway activity scores and GRN analyses, a systematic mapping was established between scFEA’s modular reaction framework and Human1 pathway annotations. This mapping was necessary because the two frameworks use different levels of metabolic granularity and nomenclature: scFEA organizes metabolism into 168 linearized reaction modules grouped into broader supermodules, whereas Human1 organizes reactions into named biological pathways. KEGG reaction identifiers were present in both frameworks and used as the bridging nomenclature.

For each scFEA module, constituent KEGG reaction IDs were matched against the rxnKEGGID field of the Human1 reactions table, establishing a many-to-many correspondence. Gene sets for matched Human1 pathways were then used to expand the scFEA gene annotations, enabling integration at the pathway level. Module-level mappings were aggregated to scFEA supermodules to support multi-resolution comparisons across all three analytical frameworks.

## Supplementary figures

**Figure S1:**
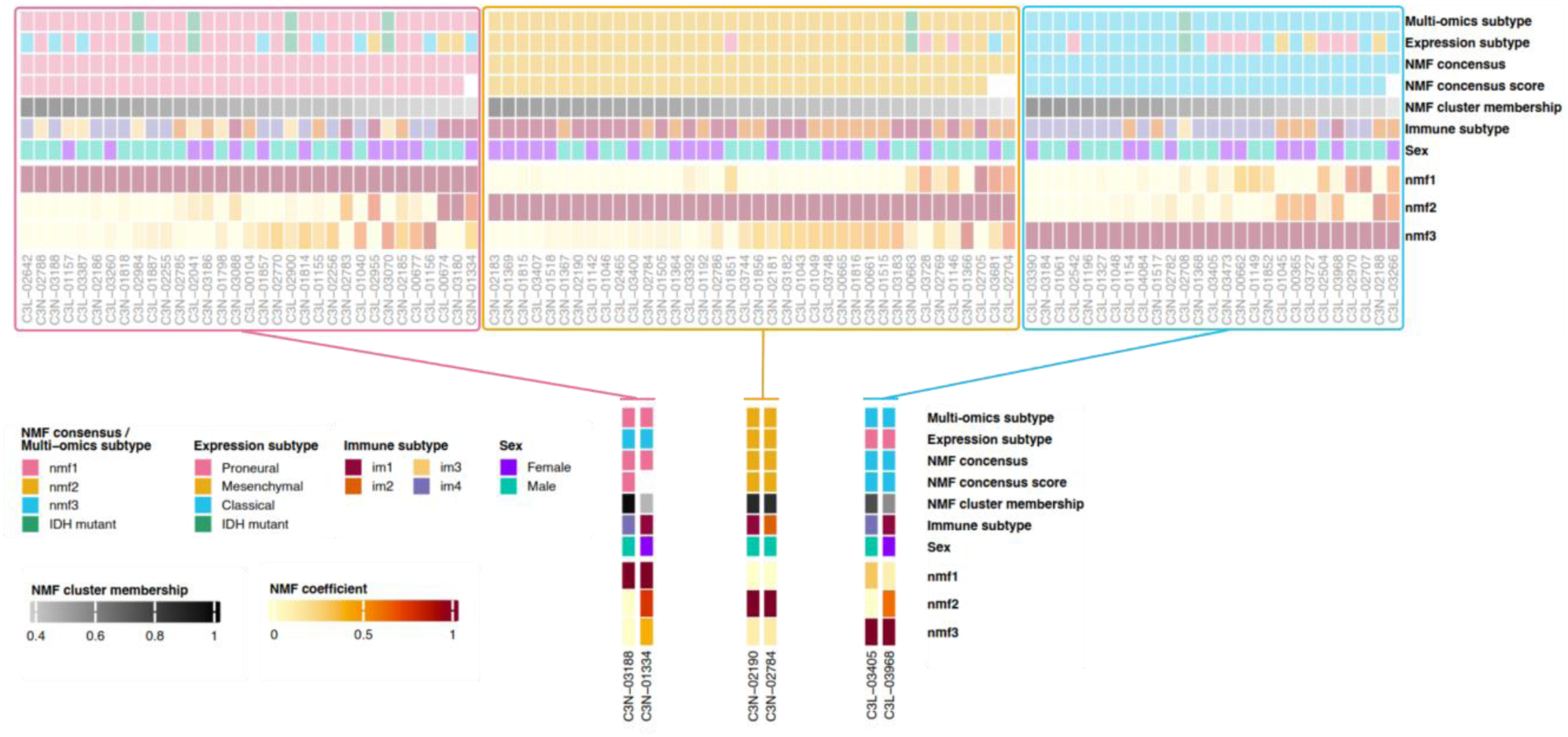
Sample overview and multi-omic subtype assessment. Heatmap of multi-omics membership scores of all three multi-omics subtypes for each tumor of the Wang dataset. Selected samples for this study are explicitly indicated

**Figure S2:**
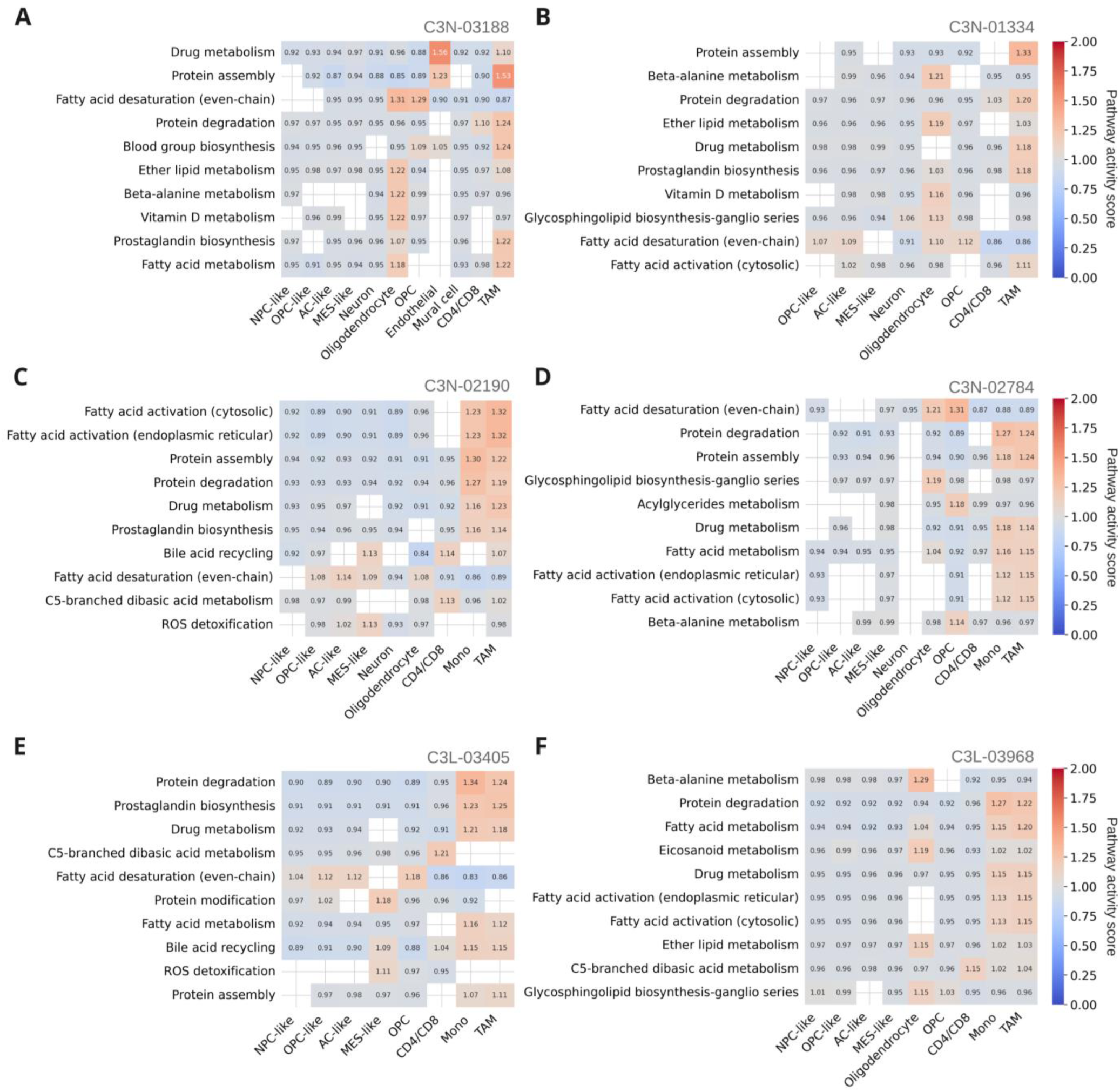
Top metabolic pathway activities for all samples. Pathway activity scores for the top 10 pathways in each of the 6 samples. Empty entries refer to non-significant activity values after permutation testing for a given pathway-cell type/state combination. Panels A and B are PN-like (nmf1) subtype samples, panels C and D are MES-like (nmf2) subtype samples, and panel E and F are CL-like (nmf3) subtype samples.

**Figure S3:**
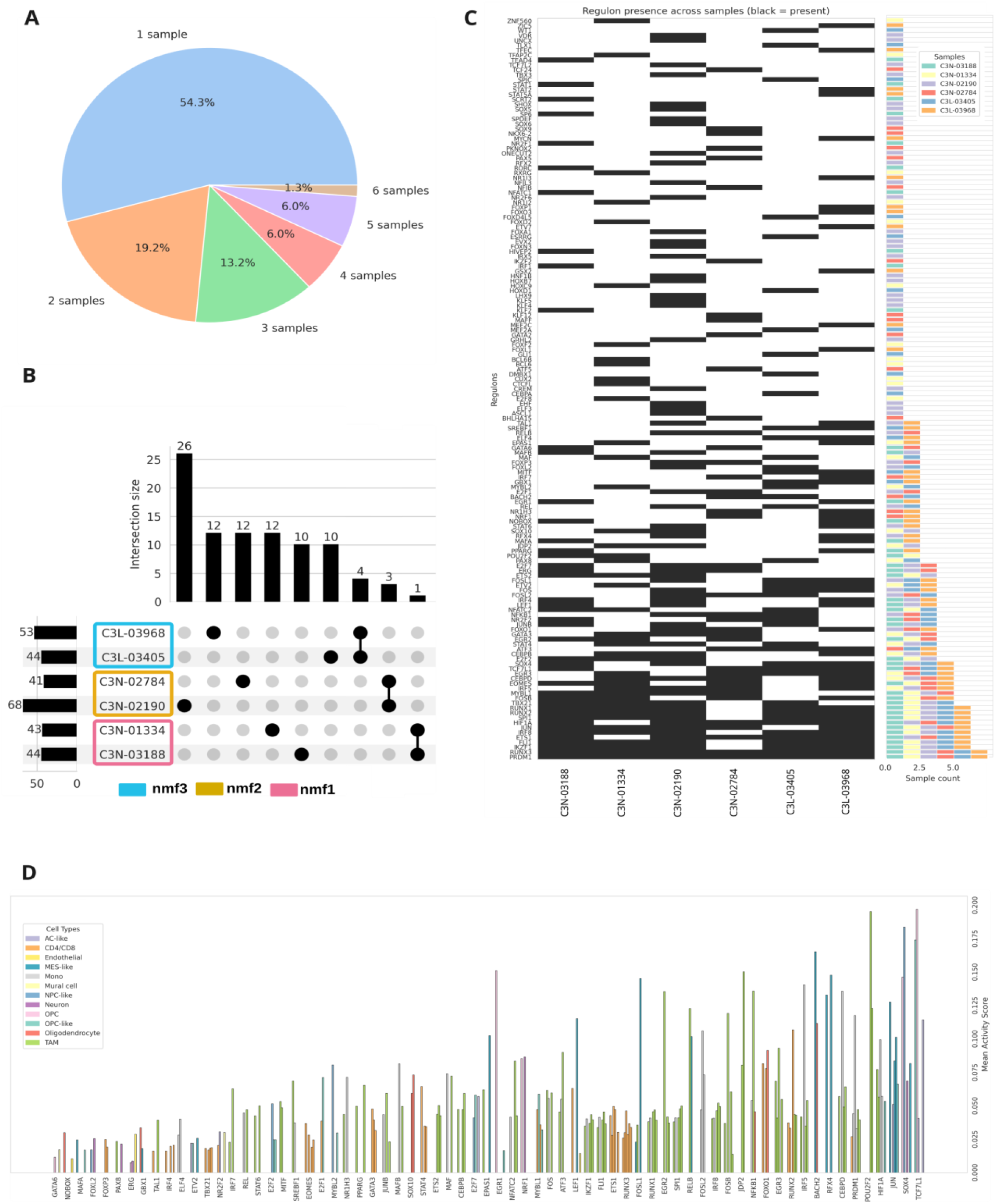
SCENIC regulon overview across samples. A) Proportion of all regulons identified in 1, 2, 3, 4, 5, or all samples. C) Binary heatmap indicating in which specific sample (columns) each regulon (rows) was identified. Black entries mean presence, white entries mean absence. B) Upset plot indicating number of sample specific regulons and shared regulons between samples of the same multi-omic subtype. D) For each regulon found in at least 2 different samples (x-axis), a bar per sample indicates in which cell type/state the regulon was highest active (color) and what the mean activity score was for that regulon in that cell type/state (y-axis).

**Figure S4:**
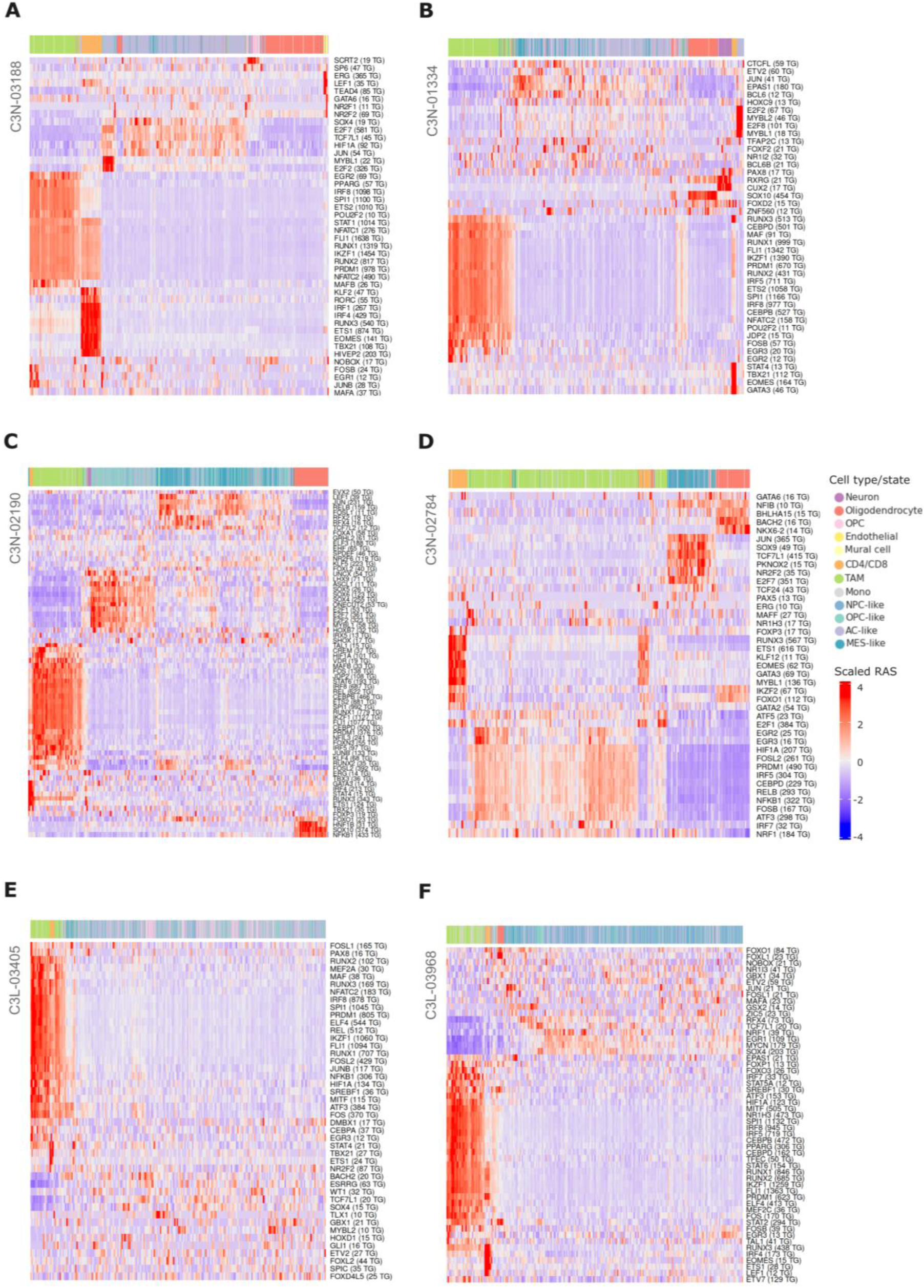
SCENIC regulon activity for all samples. Scaled regulon activity scores (RAS) across all cells for the nmf1 (PN-like) (A-B), nmf2 (MES-like) (C-D), and nmf3 (CL-like) samples (E-F). Cells are colored by cell type/state along the top color bar. Rows and columns of the heatmap are hierarchically clustered.

**Figure S5:**
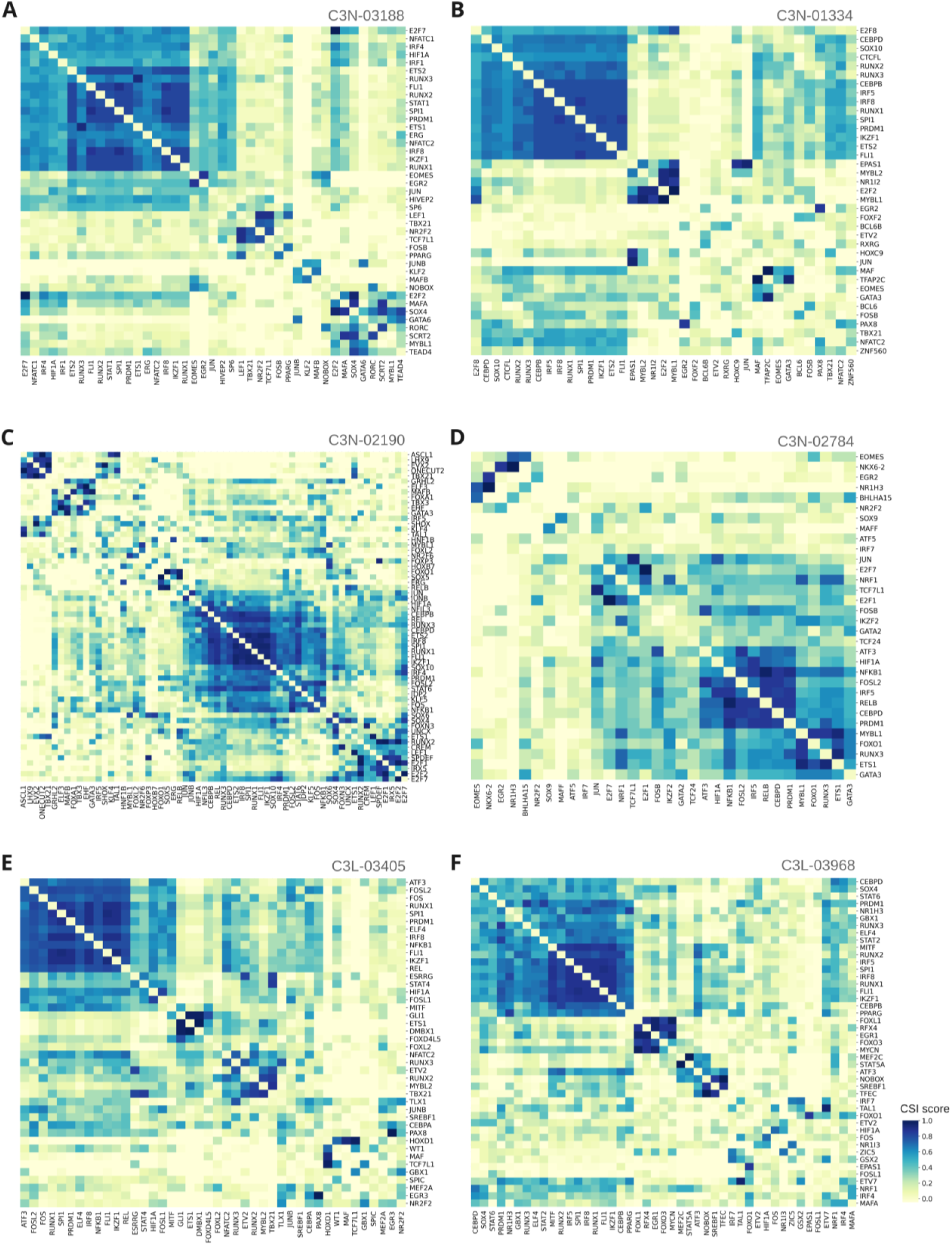
CSI scoring for all samples. CSI matrix displaying pairwise regulon similarity scores for the nmf1 (PN-like) (A-B), nmf2 (MES-like) (C-D), and nmf3 (CL-like) samples (E-F). Each entry represents the CSI score between a pair of regulons with higher values indicating more similar metabolic pathway regulation profiles. Regulons are ordered by hierarchical clustering to reveal modules of co-regulating TFs.

**Figure S6:**
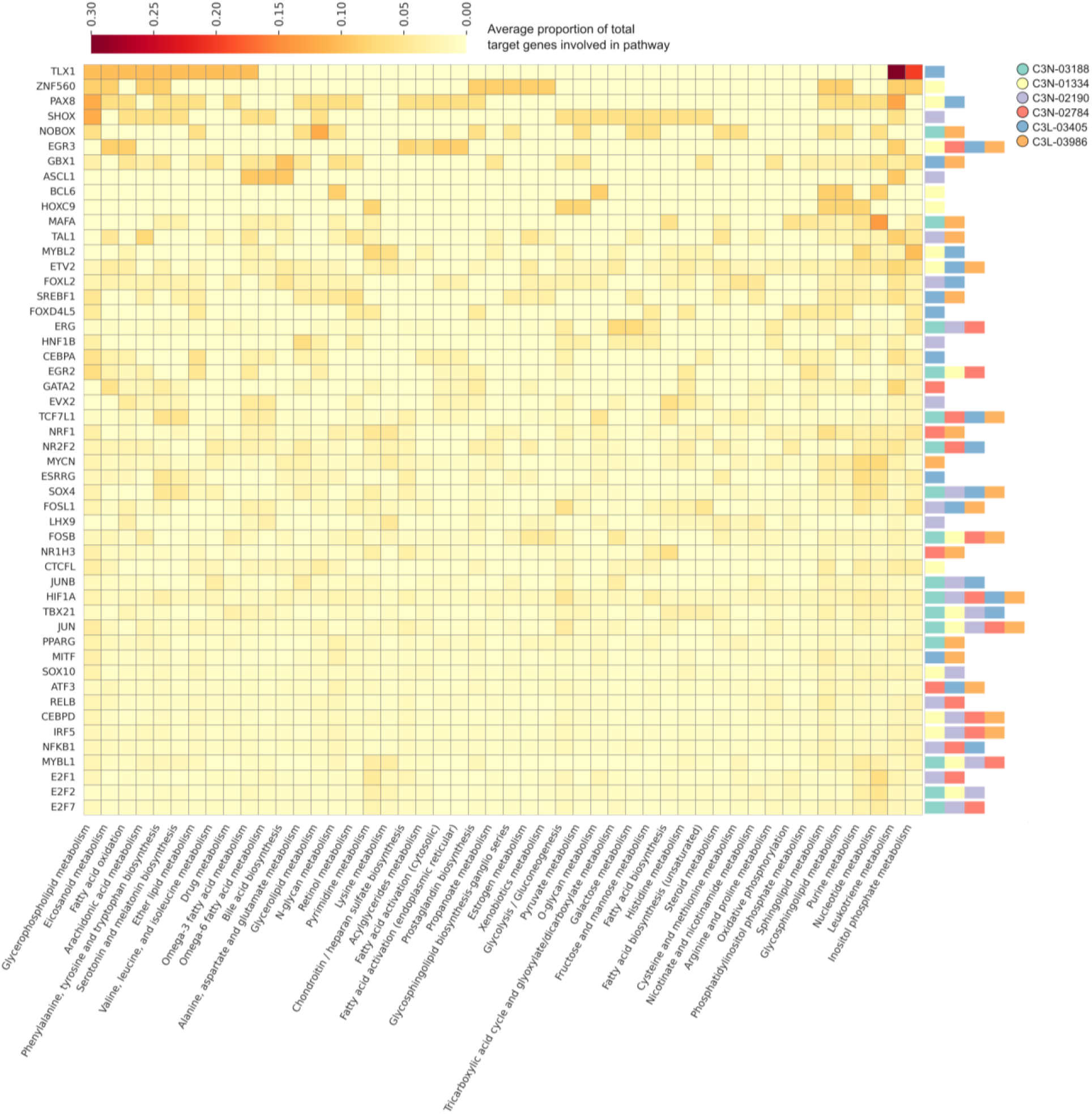
Consensus regulon x pathway overlap heatmap. Heatmap displaying the average proportion of target genes per regulon (rows) that overlap with enzyme-coding genes of Human1 metabolic pathways (columns). All zero rows and columns were removed. The stacked color bars at the right side indicate in which and how many samples the regulon was identified.

**Figure S7:**
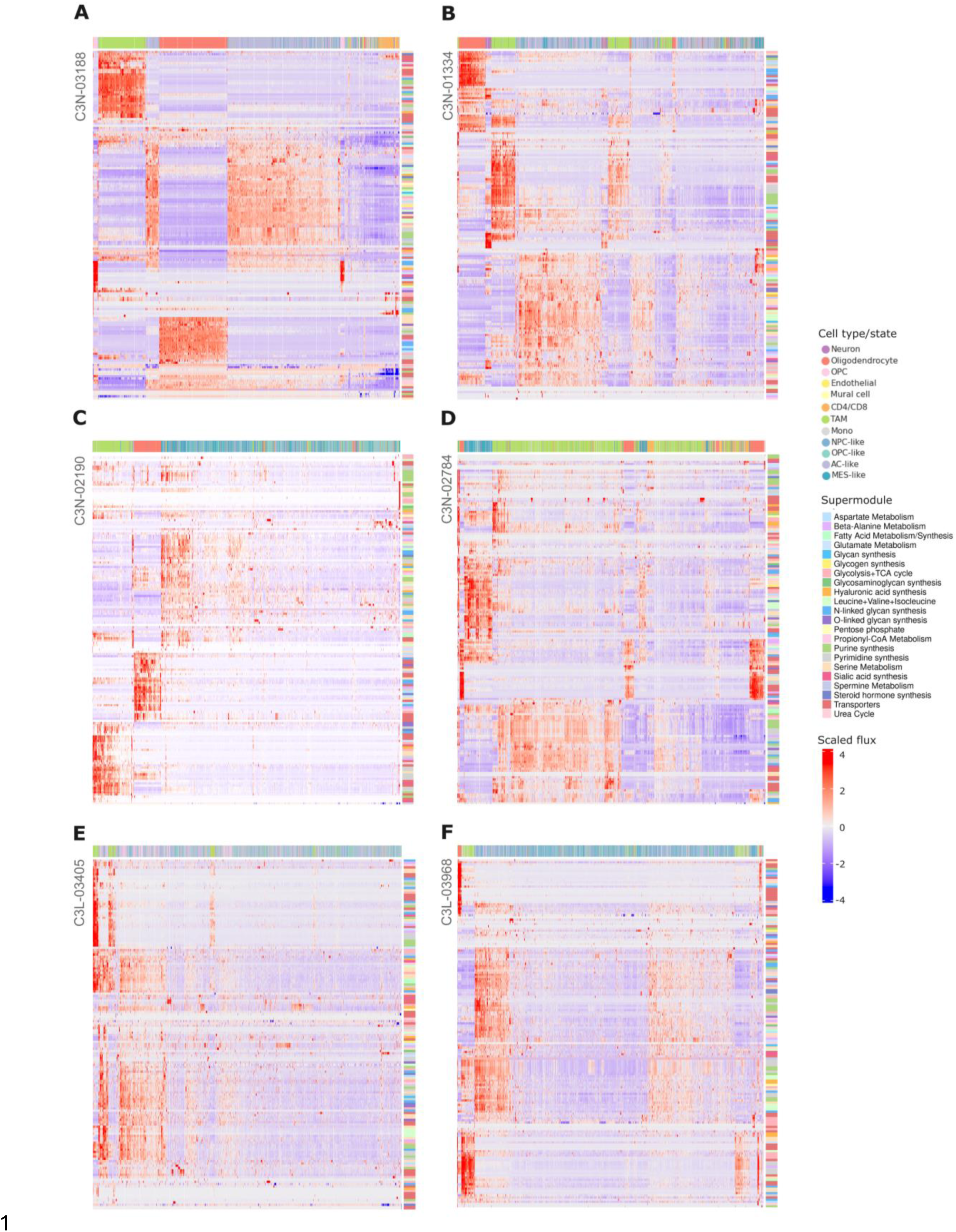
scFEA predicted metabolic flux for all samples. Heatmaps displaying scaled predicted reaction fluxes for each individual cell (columns) through each reaction module (rows) for the nmf1 (PN-like) (A-B), nmf2 (MES-like) (C-D), and nmf3 (CL-like) samples (E-F). Cells are colored by cell type/state along the top color bar. Reaction modules are colored by Supermodule on the right side. Rows and columns of the heatmap are hierarchically clustered.

**Figure S8:**
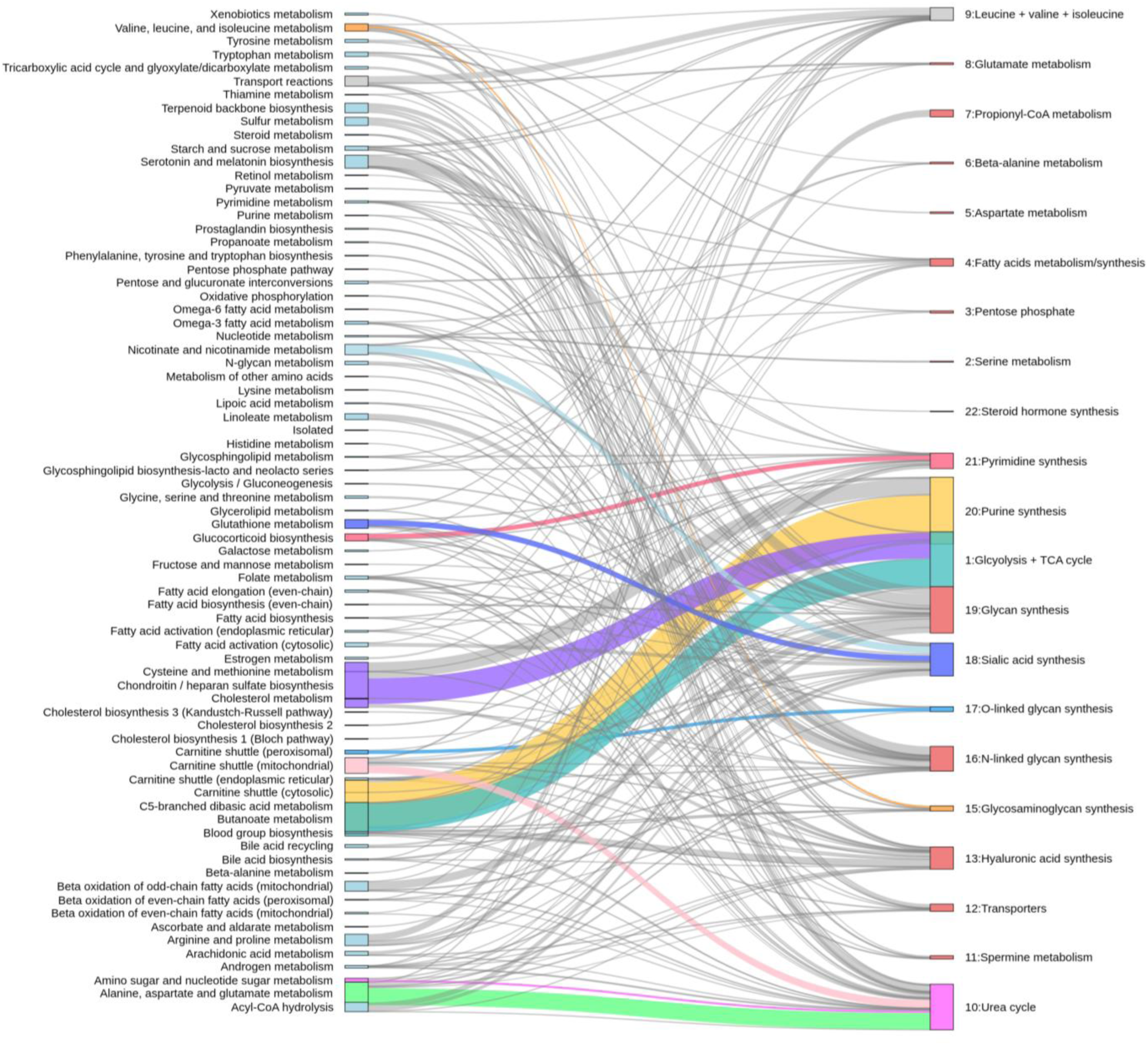
scFEA reaction module to Human1 pathway map. Individual scFEA reaction modules grouped in blocks by supermodule annotation (right) are mapped to their corresponding Human1 pathway annotation (left). The strongest Human1 pathway - scFEA supermodule pairs are colored.

